# Neural geometry of social knowledge emerging from observed social interactions

**DOI:** 10.64898/2026.02.01.703160

**Authors:** Dasom Kwon, Eshin Jolly, Luke J. Chang, Won Mok Shim

## Abstract

Humans readily infer others’ attributes and their complex web of social relationships from everyday social interactions. Yet, the neural mechanisms that transform these transient interactions into structured, multidimensional social knowledge remain unknown. Here, using a naturalistic fMRI paradigm, we develop a computational framework demonstrating how the human brain factorizes and integrates dynamic social interactions to construct multiplex social knowledge. This approach not only predicts neural responses during movie-viewing, but also enables the reconstruction of subjective social cognitive maps directly from brain activity. Crucially, the relational geometry of these reconstructed maps accurately predicts inferred personality traits, suggesting that relational and trait knowledge emerge from a shared neural representation reflecting interactional dynamics. Together, these findings provide a unifying computational framework for how the brain constructs multiplex social knowledge from observing naturalistic social interactions over time.

## Introduction

Humans have a remarkable ability to engage in complex and multifaceted social interactions (*1*). Because these interactions unfold dynamically over time (*2*), individuals must continuously track and organize information extracted from them to navigate a constantly changing social world (*3*, *4*). Social interaction constitutes the fundamental unit of the social world, through which multiple forms of social knowledge emerge (*5*), including relational knowledge, which can be formalized as a social graph that captures the structure and dynamics of interpersonal relationships (*6*).

Previous research has examined how individuals extract information from social interactions and use it to construct structured representations of relationships. For example, character networks have been modeled by quantifying the statistical *co-occurrence* of characters in comics or films, revealing graph structures that closely resemble real-world social graphs (*7–9*). Unidimensional co-occurrence-based social graphs have also been shown to explain how individuals represent and remember other people (*10*) and graph neural networks trained on co-occurrence alone can predict social evaluations without requiring complex computations (*11*). At the neural level, the perceptual presence or absence of social interactions is tracked by the lateral temporal cortex (*12*) and static social networks appear to be represented in distributed brain regions, including the medial parietal cortex and the lateral temporal and parietal cortices (*13*, *14*).

However, real-world social knowledge depends on more than simply tracking which individuals interact with one another. Rich social understanding also requires interpreting the affective and contextual dynamics through which interactions unfold over time (*15*, *16*). Existing statistical or perceptual frameworks therefore struggle to capture the inherently subjective and multidimensional structure of social interactions that give rise to social knowledge. This complexity is formally captured in network science by multiplex social graphs, which represent how the same set of individuals can be connected through multiple distinct relationships (e.g., friendships versus romances) and interaction types (e.g., fighting, laughing, teaching) (*17*). Critically, multiplex social graphs encode not only “who-interacts-with-whom,” but how individuals interact—preserving the relational and affective structure of interactions across multiple layers. Even human infants are highly sensitive to the *affective valence* of interaction, and use it to track interpersonal affiliation (*18*, *19*) that forms the building blocks of enduring relationship knowledge (*20*). In other words, dynamic patterns of positive, negative, and neutral interactions over time may provide the foundation for rich relational social cognitive representations (*21*, *22*).

Here, we propose and test whether the brain integrates observed social interactions into latent relational geometries that encode multiplex social knowledge. Using a naturalistic movie-viewing paradigm (*23*) combined with voxel-wise encoding modeling (*24*), we examined whether a quantitative model of extracted co-occurrence and affective features could predict behavioral representations of social relationships and neural responses over time. We demonstrate that social graphs reconstructed from neural activity not only reflect learned relational knowledge, but are also sufficient to accurately predict trait impressions of distinct characters. In contrast to conventional frameworks that often prioritize stable personality traits as the primary organizing principle of social knowledge (*25*, *26*), our results suggest that trait knowledge naturally emerges from the geometry of relational structures derived from observing social interactions. Together, this work introduces a computational framework for understanding how dynamic social interactions are transformed into multidimensional social knowledge and identifies the neural mechanisms that support these processes during naturalistic social experience.

## Results

Participants (N = 28) viewed a movie depicting the dynamic evolution of social relationships among six characters while undergoing functional magnetic resonance imaging (fMRI). After the movie, they rated the perceived relationship between each directed character pair across six relationship dimensions: friendship, love, trust, listening, friendship-duration, and perceived co-occurrence throughout the movie (**Fig. 1A**). Four dimensions—friendship, love, trust, and listening—were selected as the primary relationship dimensions because they capture core aspects of interpersonal affiliation and perceived relationship quality (*27*, *28*). The remaining two dimensions served auxiliary purposes: friendship duration provided a complementary temporal measure of relationship history, and perceived co-occurrence served as a behavioral validation of the computational co-occurrence features extracted from the movie. Accordingly, subsequent analyses focused primarily on the four core relationship dimensions^1^.

**Fig. 1.**
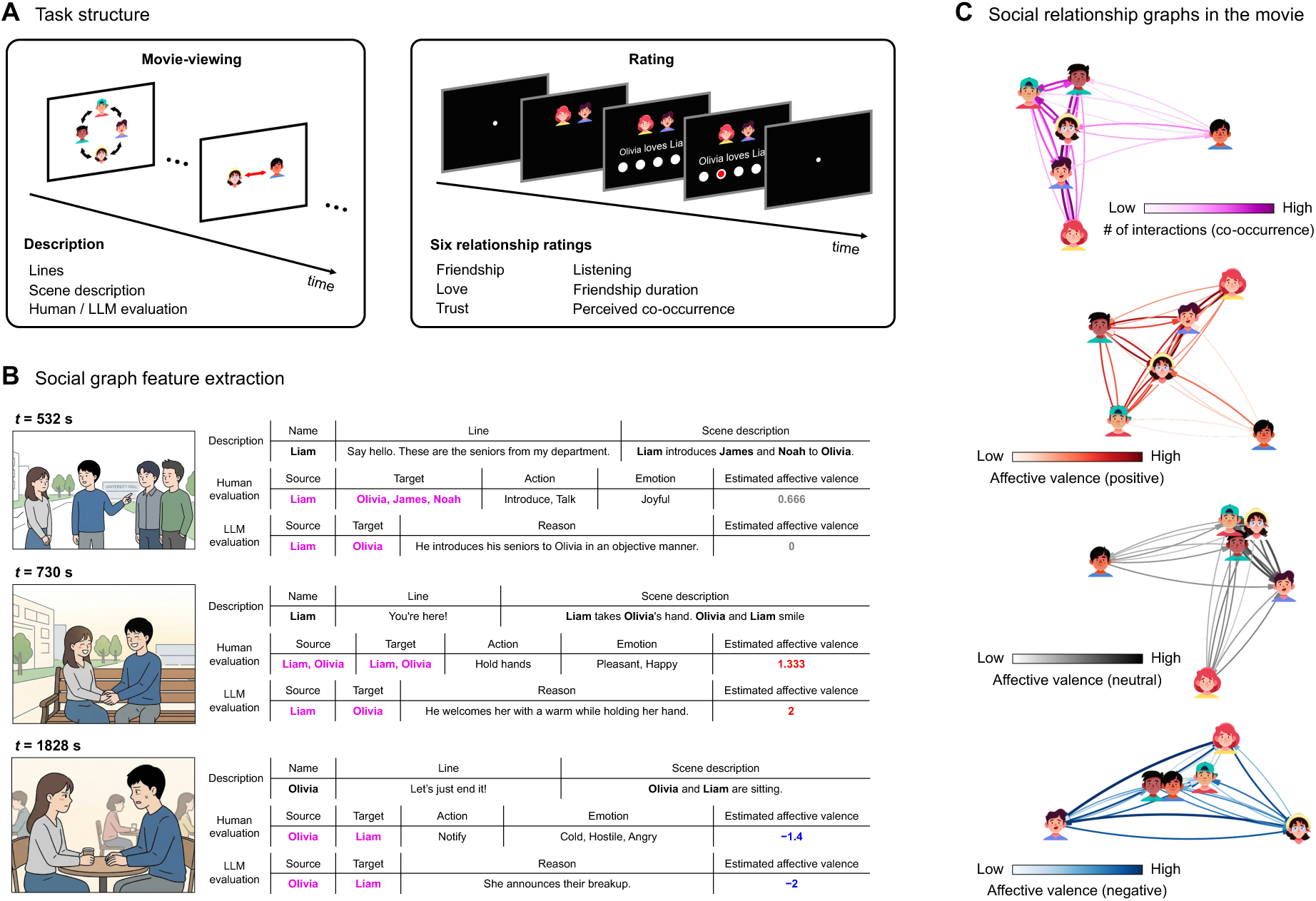
Schematic overview of the task and feature extraction. (**A**) During fMRI scanning, participants watched a movie and then rated six relationship dimensions for every directed character pair (**Table S1**). (**B**) Trained annotators provided second-by-second transcripts of characters’ lines and scene descriptions. Based on these transcripts, (1) human annotators identified the source and target characters of each interaction, along with the source character’s actions and emotions. The sentiment scores of these descriptive words were calculated using the Korean Sentiment Word Dictionary (*29*) and averaged to estimate the affective valence. (2) In a parallel approach, a large language model (LLM) was given the same descriptions as input to identify the source and target characters and estimate the affective valence. The detection of interactions showed high agreement between the human and LLM annotations (Jaccard score = 0.782). Schematic illustrations of the movie scenes were generated using Google Gemini based on the scene descriptions. (**C**) To summarize the movie narrative, event-wise features were obtained by averaging across time points within each event (as segmented by independent annotators) and subsequently aggregated. Positive, neutral, and negative interactions were quantified as independent dimensions rather than collapsed into a single valence score, preserving emotionally complex events in which opposing interaction valences co-occur. Hand-drawn face icons were used for visualization (designed by pikisuperstar on Freepik, https://www.freepik.com).

### A multiplex social graph model reconstructs relationship impressions

We first aimed to build a multiplex social graph model capturing how dynamic, evolving, affective interactions accumulate into social knowledge based on the same social signals available to participants. Building on prior work, our first graph feature captured *temporal co-occurrence* by representing characters as nodes and interactions as directed binary edges integrated over narrative events (**Fig. 1B**) (Co-occurrence_A,B_ = 1 if characters A and B appeared together) (*9–11*). Our second graph feature captured *affective dynamics* based on annotated character actions and emotions. Human annotations were converted into positive, negative, and neutral sentiment scores over directed graph edges using a pre-trained language model (*29*, *30*). These features map onto core structural properties of social network graphs, where co-occurrence establishes baseline connectivity and affective valence defines relational weighting (**Fig. 1C**). Importantly, this annotation-based approach reflects a fundamental property of social cognition: interpersonal relationships are not directly observable, but are instead inferred from the structure and affective dynamics of interactions (*15*) and therefore cannot be reduced to objective perceptual input alone (*31*). Together, these two sets of time-varying features served as input to predict participants’ subjective impressions of character relationships.

We estimated a partial least squares (PLS) regression model using leave-one-participant-out cross-validation (LOOCV). The model was trained to maximize the covariance between multiplex graph features and participants’ relationship ratings, which resulted in three latent dimensions that were sufficient to capture 95% of the variance across all rating dimensions (**Fig. 2A; see Methods**). We then evaluated model performance by computing the average cross-validated *r*^2^ across held-out participants. The multiplex social graph model explained nearly 66% of the variance in participants’ ratings (**Fig. 2B**; mean cross-validated *r*^2^ = .657, *p* < .001, interaction-shuffled permutation test). Notably, the model trained on affective valence alone outperformed the model trained on co-occurrence alone (**Fig. S1A;** mean cross-validated *r*^2^ = .601 vs. .292, feature ablation analysis). This suggests that while the perceptual detection of interactions provides a necessary baseline, observers rely more heavily on the affective quality of those interactions when forming relationship impressions. Furthermore, we evaluated the model using features automatically extracted by an LLM, which annotated interactions using only character dialogues and scene descriptions as input without any prior knowledge of the characters’ relationships. The LLM-derived features also robustly predicted participants’ relationship impressions (**Fig. S1B;** mean cross-validated *r*^2^ = .620, *p* < .001, interaction-shuffled permutation test), confirming that the model’s performance is not merely an artifact of human interpretative bias.

**Fig. 2.**
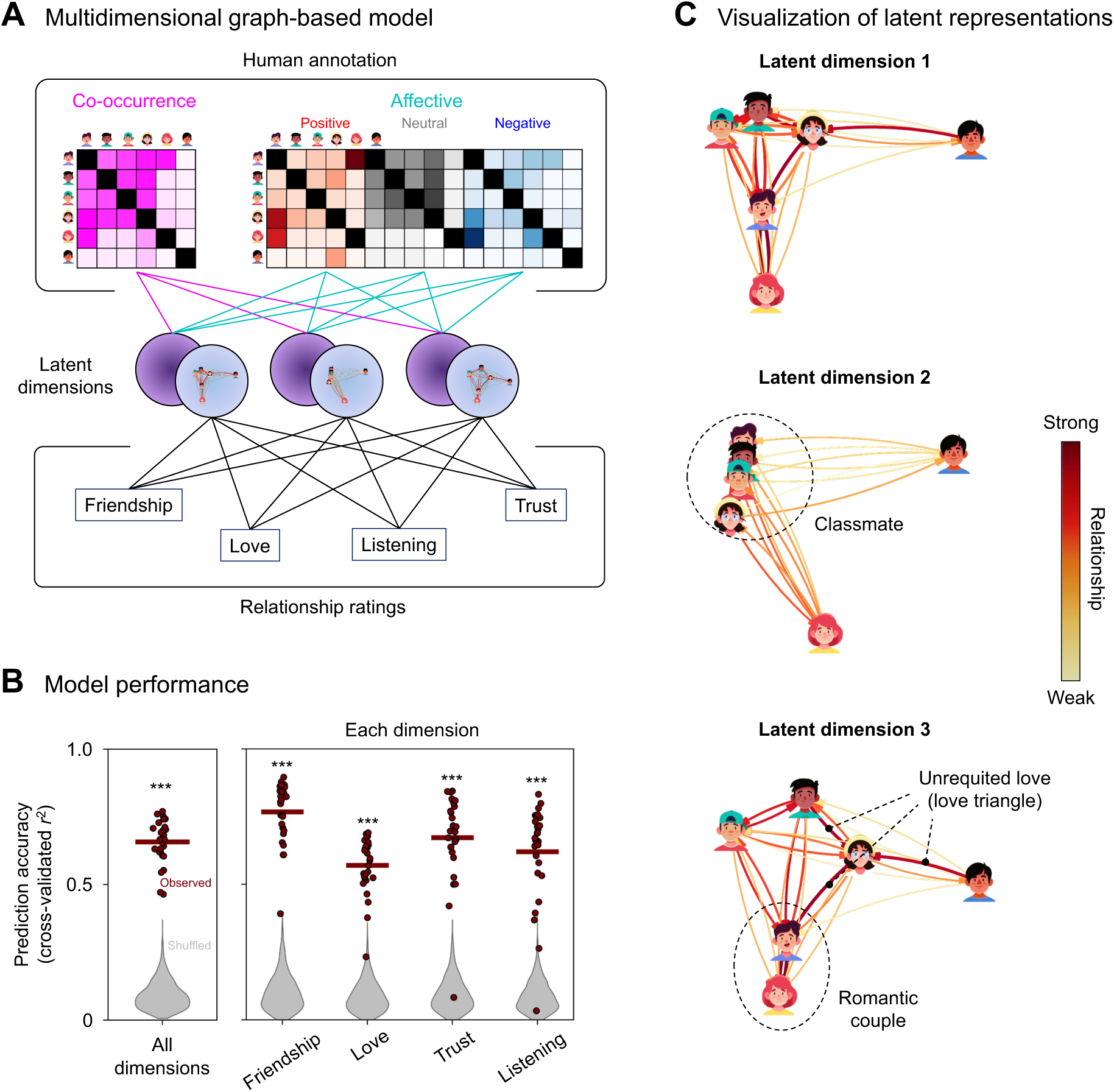
Multiplex social graph model. (**A**) Partial least squares (PLS) regression was used to simultaneously predict ratings across all four relationship dimensions from the co-occurrence and affective graphs. The number of latent dimensions was fixed at three, corresponding to the point at which 95% of the variance was explained. Predictions were evaluated using a leave-one-participant-out cross-validation scheme. (**B**) Cross-validated prediction accuracy was computed as the mean variance explained across the four relationship dimensions for each held-out participant. To generate null distributions, we randomly shuffled the identities of interacting characters in annotations at each second, repeating this process 1,000 times. (**C**) Character relationships were projected onto the model’s latent spaces, with positions estimated using multidimensional scaling. (top) The first component captures broad, symmetric cohesive ties that reflect the overall storyline. (middle) The second component reveals dense, exclusive clustering, capturing the tight-knit group of four classmates (dotted circle). (bottom) The third component captures directional, asymmetric relationship structures, reflecting complex dynamics such as unrequited love and mutual romantic bonds (dotted circle). *** *p* < .001.

Our PLS model was designed to reconstruct participants’ internal representations of their relationship impressions from interactional features. Projecting participants’ ratings onto the latent space in the model reveals how the interactional features mathematically combine to reconstruct complex relationship topologies. Rather than capturing basic positive or negative valence, the latent components captured nuanced relational structures, ranging from symmetric cohesive ties to exclusive clusters (long-term friendships among the four central classmates) and highly directional networks (romantic couples and unrequited love triangles) (**Fig. 2C, Fig. S1C**). To quantify how the reconstructed space captures structural complexity beyond affective polarity, we examined relationship asymmetry, a key feature of real-world relationships where emotional investment is often unreciprocated(*32*). We found that the topologies captured across latent components varied significantly in their degree of asymmetry; for instance, the third component exhibited significantly higher asymmetry than both the first (**Fig. S1D;** *t*(27) = 10.49, *p* < .001, two-sided paired t-test) and second components (*t*(27) = 1.98, *p* = .058).

These findings suggest that temporal co-occurrence and affective dynamics captured from social interactions are not only sufficient but constitute the computational building blocks necessary to reconstruct multiplex relational knowledge (e.g., distinguishing symmetric everyday friendships from asymmetric unrequited love).

### Multiplex social graph features predict neural activity during movie-viewing

We next examined whether multiplex social graph features not only inform complex social impressions, but also characterize neural representations that track how these social interactions unfold over time.

We used a voxel-wise encoding model approach with banded ridge regression (**see Methods**) to predict blood-oxygen-level-dependent (BOLD) responses from multiplex social graph features while participants watched the movie. Mirroring our behavioral analyses, our neural encoding model predicted each voxel’s activity at each repetition time (TR) as a linear combination of two feature subspaces: temporal co-occurrence and affective dynamics (**Fig. 3A**). We quantified the independent contributions of each subspace in explaining a voxel’s response to left-out data using six-fold cross-validation and independent hyperparameter tuning.

**Fig. 3.**
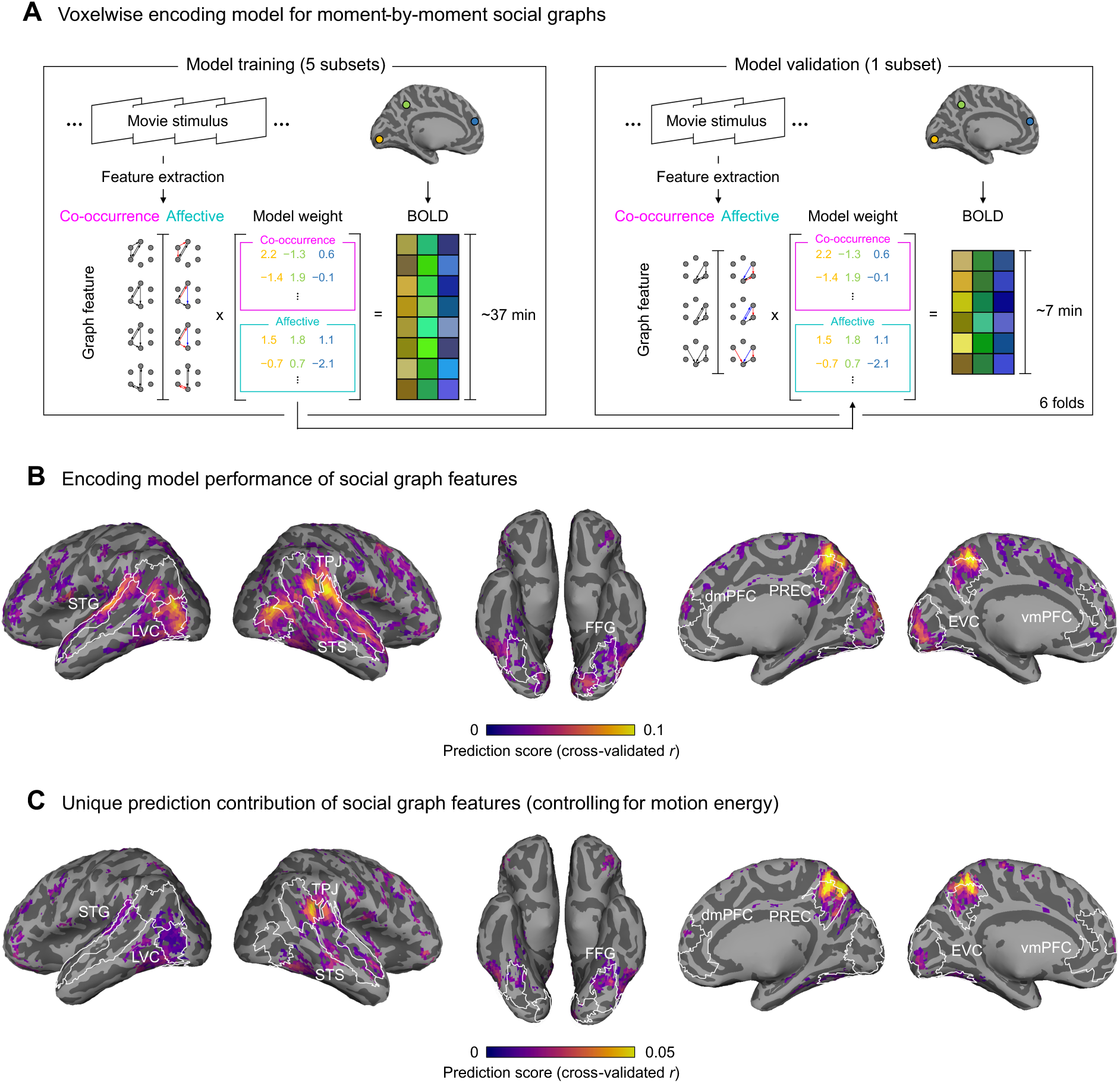
Voxelwise encoding modeling of moment-by-moment social graphs. (**A**) Movie-viewing data were divided into six folds. Model weights were optimized using five folds of training data and then tested by predicting voxelwise BOLD responses for the held-out fold, separately for each participant. Prediction accuracy was quantified as the Pearson correlation between predicted and actual BOLD responses for each voxel. (**B**) Group-level prediction accuracy maps. Only voxels with significant prediction scores are shown (*q*s < .05, FDR correction, clustering threshold = 20 voxels). (**C**) Group-level statistics of unique contributions of social graph feature controlling for low-level visual features (motion energy). Only voxels with significant unique contributions are shown (*p*s < .05, clustering threshold = 20 voxels).

The encoding model robustly predicted voxel activity across a distributed network of sensory and social brain regions. These regions included the dorsomedial prefrontal cortex (dmPFC; fraction of significant voxels: 24.66%, *q*s < .05, clustering threshold = 20 voxels, one-sided one-sample bootstrap test), ventromedial prefrontal cortex (vmPFC; 8.53%), precuneus (PREC; 67.57%), temporoparietal junction (TPJ; 45.55%), superior temporal sulcus (STS; 57.09%), superior temporal gyrus (STG; 80.29%), fusiform gyrus (FFG; 35.57%), lateral visual cortex (LVC; 84.35%), and early visual cortex (EVC; 43.19%) (**Fig. 3B**; **Fig. S2**). We further confirmed the multiplex social graph model’s robustness by testing additional variants that controlled for low-level visual signals (e.g., motion energy) (**Fig. 3C**) or considered alternative timescales of dynamic interactions (**Fig. S3**).

Furthermore, each multiplex graph feature subspace uniquely contributed to predicting voxelwise BOLD responses (**Fig. S4**). Co-occurrence features better predicted activity in sensory cortices, including the ventral visual stream and the lateral part of the sensory brains (**Fig. S5**), while affective features more strongly recruited the TPJ, PREC, and mPFC. These findings are consistent with previous work demonstrating a hierarchical organization of social action features along the lateral visual pathway (*33*) (i.e., detecting the presence of interactions from visual cues) and social-affective features encoded in cortical midline regions and transmodal brain areas (*34*, *35*).

### Distributed cortical representations of relational topologies

While the previous analysis demonstrated how multiplex social graph features can reliably predict brain activity over time, we next asked whether these learned voxel feature weights jointly yielded interpretable representations that could be used to predict participants’ relationship impressions. In other words, we examined whether the latent geometry extracted from neural responses captures the subjective cognitive map underlying participants’ evaluation of social relationships.

To do so, we adapted the same PLS regression approach from our behavioral analysis, but this time using learned voxel feature weights as inputs to predict participants’ relationship ratings (**Fig. 4**). Because learned weights for each encoding feature correspond to each directional pairwise-edge of the co-occurrence and affective multiplex graphs, we can use these weights as a basis set to reconstruct participants’ subjective impressions of character relationships, akin to inverted encoding models (IEM) used to reconstruct stimuli from neural population responses (*24*, *36*). We quantified cross-validated reconstruction accuracy using the Pearson correlation between predicted and actual relationship ratings.

**Fig. 4.**
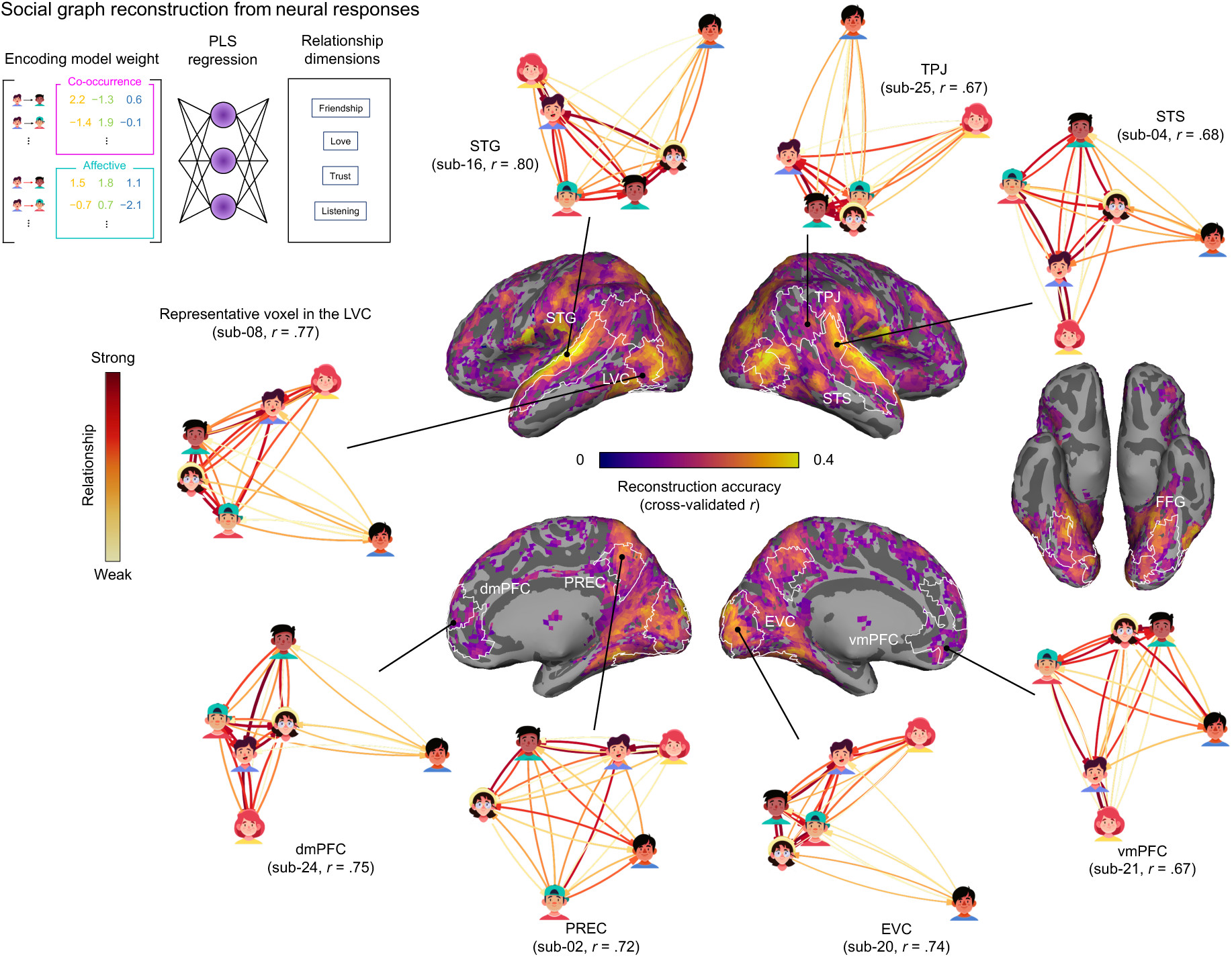
Reconstruction of participants’ relationship impressions using learned multiplex model features. Social relationship graphs were reconstructed from voxelwise encoding model weights using PLS regression with a LOOCV scheme. Reconstruction accuracy was quantified as the Pearson correlation between the predicted and actual four-dimensional relationship ratings. Only voxels with significant reconstruction accuracy are shown (*q*s < .05, FDR correction, clustering threshold = 20 voxels). The visualized graphs show the weighted sum of the X-side latent representations (neural representations), combined according to their explained variance ratios, reconstructed from voxels within representative brain regions for individual participants. Each reconstructed graph is accompanied by labels indicating the brain region, participant ID, and reconstruction accuracy.

As seen in **Fig. 4**, multiplex feature weights successfully reconstructed ratings across a wide range of cortical regions, spanning both sensory and higher-order areas (*q*s < .05, clustering threshold = 20 voxels, two-sided one-sample t-test). Importantly, the successful reconstruction in key social brain regions (e.g., mPFC, precuneus) could not be explained by low-level stimulus properties, as controlling for visual features selectively reduced decoding performance in lateral visual areas while leaving the higher-order representational maps intact (**Fig. S5A**). However, as visually highlighted, reconstructed social graphs varied substantially across voxels and participants, obscuring the richness and complexity that each region was actually encoding. To unpack this complexity, we identified the minimum number of latent components required for each voxel-wise encoding model to achieve 95% of its maximum reconstruction accuracy. This confirmed the heterogeneity of learned representations (**Fig. S5B**): some regions encoded simple, low-dimensional aspects of relationships, while others captured richer, multiplex structures. At the same time, relationship impressions of characters varied across participants and relationship dimensions (e.g., how similarly participants perceived friendship-based (ISC *r* = .93) compared to romantic relationships (ISC *r* = .62)) (**Fig. S6**).

In other words, unlike recent advances in visual and linguistic neural encoding models that have a clearly identifiable objective outcome shared across participants (i.e., words in a story, images on a screen) (*37*), social relationships are the product of both exogenous (stimulus) and endogenous (personal, subjective) aspects of social inferences (*38*). However, individuals still experience measurable alignment in their mnemonic and neural responses when entraining to the same naturalistic stimuli (*10*, *39*). This raises the question: Can we leverage inter-individual alignment to discover shared multiplex social graph representations across participants?

### Shared low-dimensional geometry of social relationship representations

Because our neural encoding models quantify the contribution of the same multiplex graph features across participants (i.e., same co-occurrence and affective information), we estimated a Shared Response Model (SRM; **Fig. 5A; see Methods**) to identify a reduced space of encoding features common across all participants and voxels. Using SRM in this way leverages intersubject correlation (ISC) as a content-agnostic measure (*40*) to jointly learn shared multiplex graph representations across regions and individuals. Rather than asking whether a single pair of participants shared similar neural encoding features in a specific region, we asked whether all participants share common multi-voxel patterns that jointly encode their shared impressions of social relationships.

**Fig. 5.**
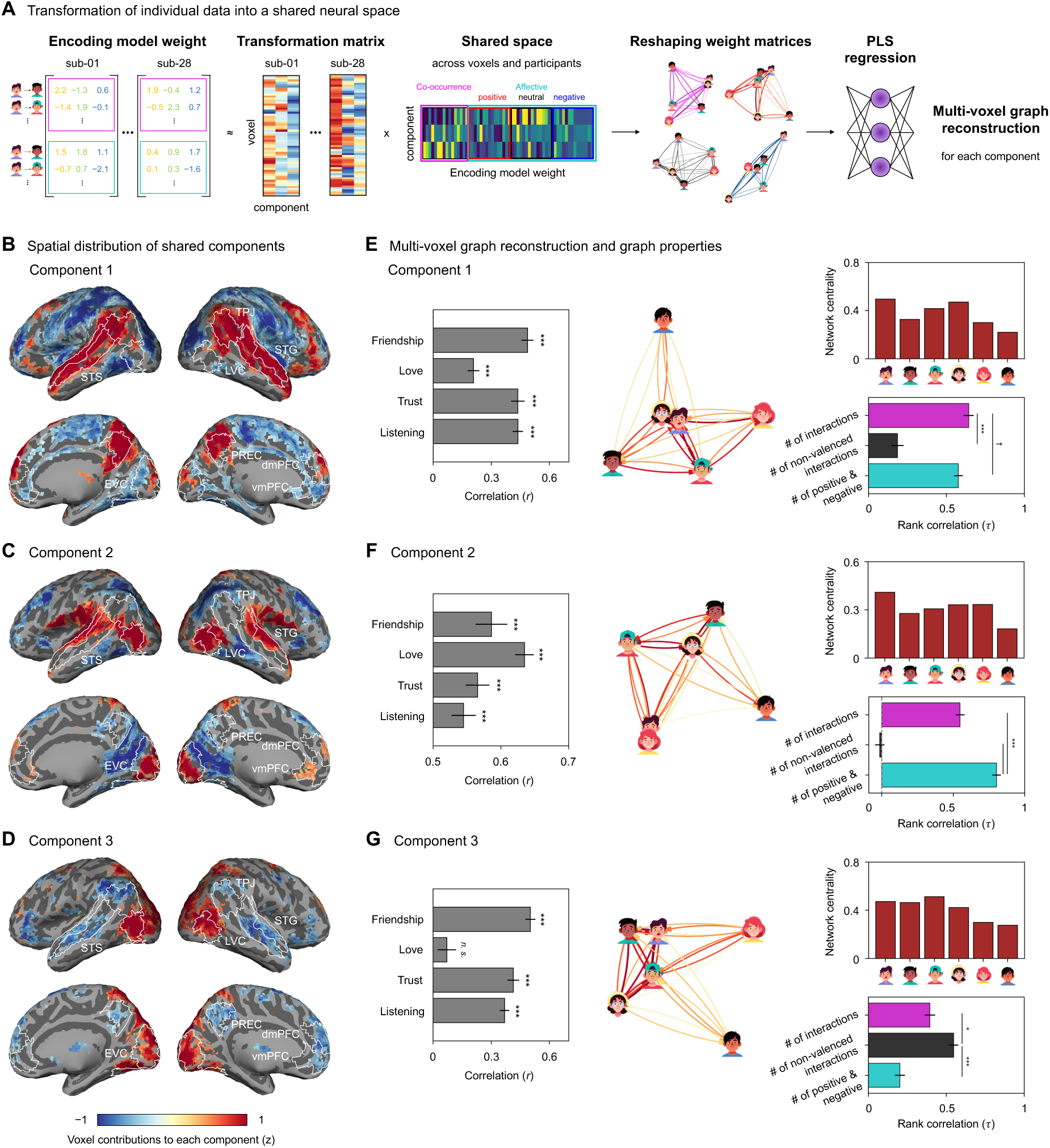
Shared neural space for social graph encoding. (**A**) Encoding model weights from 28 participants were transformed into a shared neural weight space using the Shared Response Model (SRM). The shared neural weights were then reshaped into four graph types (co-occurrence and three affective graphs) and used to predict participants’ four-dimensional relationship ratings. (**B-D**) Each brain map depicts voxelwise loadings from the SRM transformation matrices across participants. Voxel loadings were averaged across participants to visualize consistent spatial patterns (*q*s < .05, FDR correction, clustering threshold = 20 voxels). (**E-G**) Cross-validated reconstruction accuracy was computed as the Pearson correlation between predicted and actual responses across the four relationship dimensions for each held-out participant. The visualized graphs show the weighted sum of the X-side latent representations, combined according to their explained variance ratios. To interpret the neural representations, we computed node-level network-centrality for each character within each shared component and compared these centrality profiles to those derived from annotation graphs quantified by different interaction types. Component 1 showed the highest similarity with networks based on total interaction frequency. Component 2 corresponded to networks weighted by positive and negative valenced interactions, and Component 3 aligned with networks reflecting non-valenced interactions. It is important to note that SRM loadings indicate the direction of contribution within the shared representational space rather than the magnitude of neural responses. † *p* = .071, * *p* < .05, *** *p* < .001.

Using LOOCV voxelwise reconstruction accuracy as our principled selection criterion for choosing the number of SRM components, we learned a solution with three stable latent components (ISC *r*: Component 1 = .84, Component 2 = .90, Component 3 = .88) (**Fig. S7**). Moreover, each component reflected distinct contributions from interpretable large-scale cortical networks (**Fig. 5**). Importantly, the SRM does not restrict latent components to be orthogonal or independent, so individual voxels contribute to every SRM component in different ways. The first component comprised alignment in temporal cortices, PREC, and dmPFC (**Fig. 5B**). The second component was characterized by aligned contributions from sensory cortices and mPFC (**Fig. 5C**). The third component consisted of alignment spanning from sensory cortices to parietal regions (**Fig. 5D**).

We then repeated our PLS regression approach as before, but this time using SRM components reshaped to represent the original pairwise multiplex social graph features used to train the neural encoding models (**Fig. 5A**). This approach allowed us to directly examine shared neural representations of relational geometries across participants. Specifically, we estimated a separate PLS regression for each SRM component to predict participants’ relationship ratings, and quantified reconstruction accuracy using LOOCV. The social graph reconstructed from the first SRM component primarily represented participants’ impressions of friendship (**Fig. 5E**), while the second SRM component primarily reflected impressions of the love dimension (**Fig. 5F**). The third SRM component’s graph reconstruction resembled that of the first (**Fig. 5G**). To quantitatively compare these reconstructions, we computed the network centrality of each character and compared it to annotated features originally used to construct our multiplex social graph models: (1) the total number of interactions, (2) the number of non-valenced interactions, or (3) the number of emotionally valenced interactions (**Fig. 5E-G**).

This analysis clearly revealed distinct contributions of each SRM component despite superficially similar reconstructed social graphs. Node centrality within the first SRM component most closely tracked interaction frequency (**Fig. 5E**), whereas centrality within the second SRM component was primarily driven by affectively weighted frequency (positive or negative interactions specifically) (**Fig. 5F)**. Notably, despite reconstruction similarity with the first component, node centrality within the third SRM was markedly different: it reflected information from non-affective interactions (**Fig. 5G)**. In other words, shared graph representations across participants simultaneously captured distinct aspects of the relationships between characters. Because voxels contribute to every SRM component, our approach disentangles the multiplexed graph features that inform participants’ relationship impressions. Each SRM component can be interpreted as a different “view” of the social graph informed by differential contributions of co-occurrence and affective features. Unlike our previous analysis that independently reconstructed a separate graph per voxel and participant, this analysis leverages the shared information across both voxels and participants to discover the most stable views (layers) of the multiplex social graph that reliably capture distinct properties of individual characters (nodes).

### Relational graph representations predict trait impressions

Motivated by the previous result identifying different “perspectives” of each character’s centrality (within each SRM component), we asked: Can shared graph representations also predict richer character impressions like *personality traits*? This question is especially intriguing given that these two relational and person-level knowledge have largely been examined independently (*3*, *41*). To explore this, we calculated several additional node-level graph metrics for each character separately within each SRM component, reflecting relational features at the node level: edge strength (the mean of in-going and out-going), edge strength ratio, reciprocity, and local clustering coefficient (**Fig. 6**). We then used all features across all components (3 SRM components x 4 metrics = 12 total) to predict participants’ trait impressions for each character (dominance, warmth, extraversion, competence, trustworthiness, openness, and attractiveness) using ridge regression with LOOCV and quantified prediction accuracy using Pearson correlation.

**Fig. 6.**
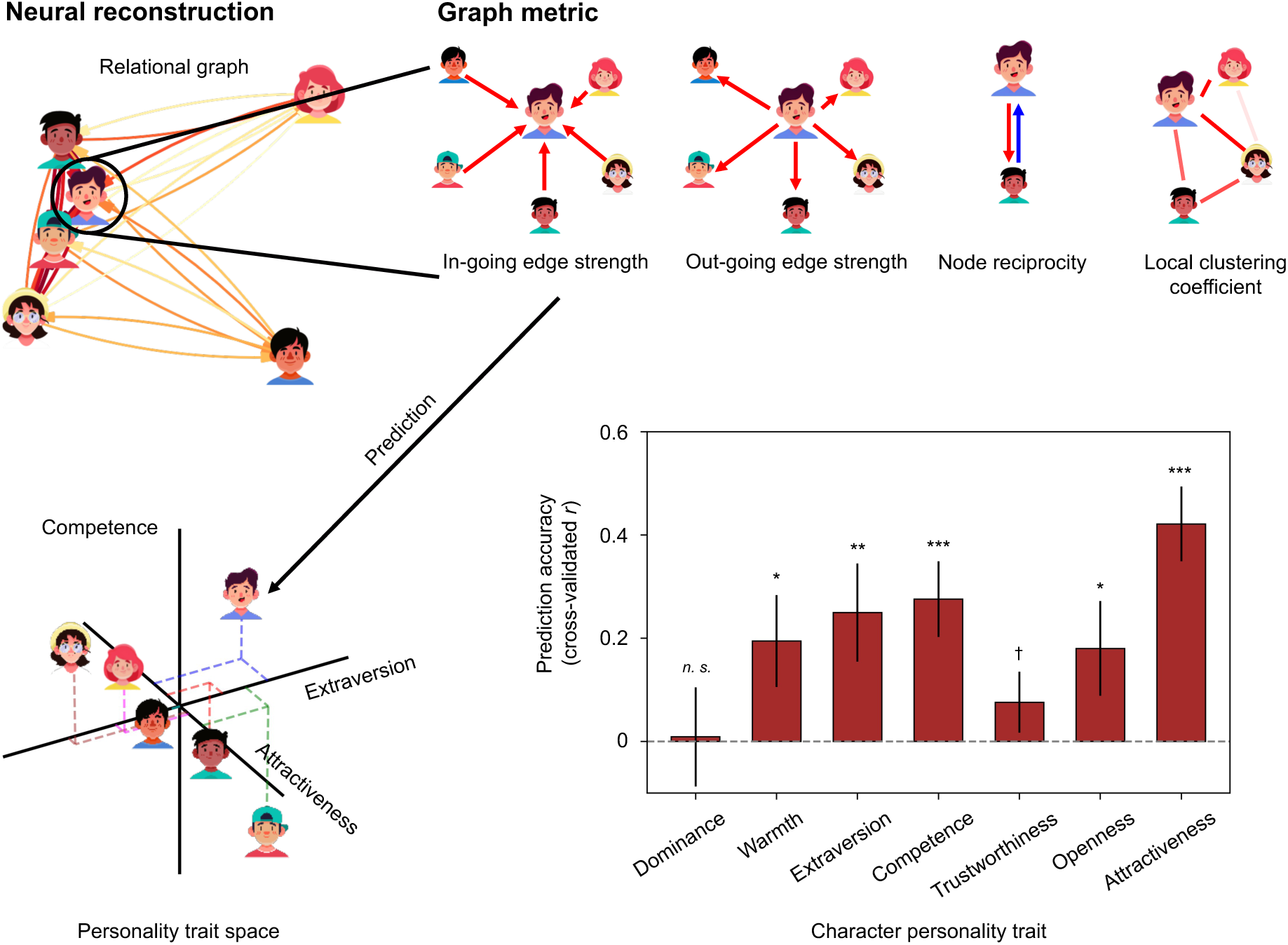
Personal knowledge within interaction-based social relationship graphs. Node-level graph metrics for each character were extracted from the neural reconstruction of social relationship graphs, including in-going and out-going edge strength, in-out edge strength ratio, node reciprocity, and local clustering coefficient. These graph-derived metrics from the three shared components were then used as predictors in a ridge regression model to estimate seven personality trait ratings (dominance, warmth, extraversion, competence, trustworthiness, openness, and attractiveness) reported by participants. † *p* = .096, * *p* < .05, ** *p* < .01, *** *p* < .001.

Despite no information being provided to our multiplex social graph model, neural encoding models, or SRM analyses about participants’ trait impressions, we significantly predicted participants’ impressions of warmth, extraversion, competence, openness, and attractiveness (cross-validated *r*s > .18, *p*s < .05, one-sided one-sample t-test) (**Fig. 6**). The fact that some trait dimensions were not reliably predicted may reflect stimulus-specific limitations rather than a general limitation of the model. In particular, dominance was not significantly predicted, likely reflecting limited trait-relevant information in the movie stimulus used in this study, leading to increased inter-individual variability in dominance judgments. Consistent with this interpretation, dominance ratings showed the lowest agreement across participants (ISC *r* = .001; for other traits, ISC *r*s > .088), indicating little shared information across participants. These results demonstrate that multiplex relational graph representations are sufficient to support accurate predictions of trait impressions from interaction patterns, without any explicit trait information. This capacity to predict trait impressions from relational structure alone suggests that impressions of others’ traits naturally emerge from observing social interactions, bridging relational and person-level social judgments within a shared representational framework.

## Discussion

In this study, we developed a computational framework to model how the brain encodes dynamic social interactions using naturalistic stimuli. Our results show that the brain constructs multiplex social knowledge through interaction-based representations that integrate temporal co-occurrence structure with affective dynamics. Specifically, our multiplex graph model not only predicted neural responses during movie viewing but also enabled the high-fidelity reconstruction of subjective cognitive maps directly from brain activity. Furthermore, the relational geometry of these reconstructed maps successfully predicted individual node-level information, including inferred personality traits. Rather than conceptualizing relational and person-level knowledge as distinct forms of social understanding, our results suggest that both emerge from a shared representational basis grounded in observed social interactions. Together, these findings highlight that social interactions serve as the primary organizing units that are integrated over time to form multiplex representations in the brain.

Social interactions convey both statistical and affective cues that unfold dynamically in naturalistic settings (*33*, *42*). For social understanding, the brain must integrate information about who interacts with whom with inferences about the internal states that give meaning to those interactions (*34*, *43*). Our findings suggest that the cortex factorizes moment-by-moment interactions into separable but jointly interpretable subspaces: temporal co-occurrence and affective dynamics. At a computational level, tracking co-occurrence relies on low-level statistical operations that monitor how often entities appear together (*8*, *44*, *45*), consistent with processing streams in sensory cortices. Conversely, interpreting affective dynamics requires inferring others’ internal states (*46*, *47*), which aligns with computations typically associated with core mentalizing regions. Under this computational account, multiplex graph features are not merely descriptive annotations of the narrative, but representational primitives that the brain can integrate over time to organize higher-order relationship structures (*48*).

The feature weights derived from the voxel-wise encoding model can be interpreted as neural tuning functions for observing moment-by-moment social interactions. By reconstructing the global relational geometries among characters from these learned weights, we show that macroscopic social knowledge is constructed through the integration of these microscopic tuning profiles. This process parallels computational frameworks in other cognitive domains: just as orientation tuning functions form the basis for complex visual representations (*49*, *50*), and spatial tuning functions underlie cognitive maps of the physical environment (*51*, *52*), our findings suggest that the brain utilizes interaction tuning functions as the geometric basis for constructing multidimensional social knowledge. This framework formally bridges the transient perception of social interactions (*33*, *53*) and the encoding of stable social networks (*13*, *54*). Finally, the emergence of multiple shared dimensions across individuals suggests that social relational knowledge is organized as a set of interaction axes, each capturing a different facet of social relationships in the world. These dimensions may also reflect distinct modes of naturalistic stimulus processing, such as social attention or affective evaluation (*55*, *56*). Rather than being mutually exclusive interpretations, we suggest that these components may reflect the specific large-scale brain networks (*57*, *58*) recruited to integrate different layers of relational knowledge from dynamic interactions.

Previous studies in social neuroscience have made significant strides in mapping the organization of social knowledge, often demonstrating that person-level knowledge (e.g., traits and mental states) and relational structures (e.g., social networks) are represented within a structured space (*3*, *14*, *59*). The present findings extend this perspective by proposing that both forms of social knowledge may arise from a common interaction-based foundation, framing them as complementary projections of a shared underlying interaction history. By emphasizing moment-by-moment social interactions, our results illustrate how higher-order social knowledge emerges from dynamic interaction patterns (*60*). Crucially, this implies that interaction-derived relationship representations may give rise to person-specific knowledge. Rather than viewing individuals as collections of attributes inferred independently of their relational context (*26*, *61*), trait impressions emerge from how individuals are embedded within a multiplex social graph. Under this account, impressions of personality are constructed by integrating patterns of interaction and their affective qualities over time, analogous to how temporal sequences are organized into latent representations of space (*62*). Together, these findings suggest that the tightly interconnected space of individual and relational knowledge is fundamentally grounded in the continuous observation of social interactions.

While our model captures shared relational geometry across participants, relationship inference is inherently subjective. Unlike perceptual domains where a shared external referent can serve as an approximation of ground truth, social relationships have no single objective ground truth (*15*, *63*)—they are jointly constructed by the observable structure of interactions and the interpretive processes of individual perceivers. This is not a limitation of the present approach but a fundamental property of social cognition itself, and one that our framework explicitly embraces by modeling the shared structure that emerges across perceivers while preserving the subjective variability inherent to social inference (*38*, *57*). Future work should investigate how specific perceiver characteristics, such as observers’ own personality traits, self-referential anchoring tendencies, or their perceived social distance from the characters, modulate the neural representations underlying social inference (*64*). Understanding how individual differences in observers’ idiosyncratic interpretations shape their perception of dynamic interactions will provide a more complete account of how the human brain constructs individualized social cognitive maps (*65*).

Building on this interaction-based account of social knowledge, our results suggest a representational organization in which relational knowledge is structured along interpretable interaction axes that capture regularities in how individuals engage with one another over time. This interaction-based organization parallels cognitive map frameworks, in which continuous experience transforms into structured representations that support flexible inference and generalization (*21*, *66*, *67*). Extending these frameworks to the social domain implies that social knowledge is represented within a structured space defined by distinct interaction axes. While prior work has similarly shown that social information occupies low-dimensional spaces (*41*), our findings ground such abstract dimensions within an interaction-based framework, demonstrating that they reflect the accumulation of social interactions over time.

## Acknowledgments

This work was supported by the National Research Foundation of Korea and the Ministry of Education of the Republic of Korea (RS-2024-00348130, RS-2025-02304581, NRF-2025S1A5A2A03013734), and the Fourth Stage of Brain Korea 21 Project in the Department of Intelligent Precision Healthcare, Sungkyunkwan University (S-2023-0794-000) to W.M.S.

D.K. was supported by funding from the Institute for Basic Science (IBS-R015-D2).

## Author contributions

**D.K.**: Conceptualization, Methodology, Formal analysis, Investigation, Data curation, Visualization, Writing - Original Draft, Writing - Review & Editing.

**E.J.**: Conceptualization, Methodology, Formal analysis, Writing - Review & Editing.

**L.J.C.**: Conceptualization, Methodology, Formal analysis, Writing - Review & Editing.

**W.M.S.**: Conceptualization, Methodology, Formal analysis, Writing - Review & Editing, Funding acquisition

## Declaration of Interests

The authors declare no competing interests.

## Data availability

The behavioral and fMRI data are currently available for reviewers at https://drive.google.com/drive/folders/17W6mkoSVG6KdKL8dIoZXd6zUnLJid2TD?usp=drive_link and will be deposited in OpenNeuro upon publication.

## Code availability

Analysis codes are available at https://github.com/somvid/Dynamic-social-interactions.

## Methods

### Participants

Twenty-eight participants aged 19 to 35 years (*M* = 23, *SD* = 3.16, 12 females) were recruited from the Sungkyunkwan University community. All participants were fluent in Korean, had normal or corrected-to-normal vision, and normal hearing. They received monetary compensation for participating in a functional magnetic resonance imaging (fMRI) experiment. Prior to the study, they provided informed consent in accordance with the guidelines of the Institutional Review Board of Sungkyunkwan University.

### Task structure

All tasks were performed inside an fMRI scanner (**Fig. 1A**) and were presented using Matlab with Psychophysics Toolbox (*68*).

#### Movie-viewing task

Participants watched the first season of the Korean web drama *Love Playlist* (available [online](https://www.youtube.com/playlist?%20app=desktop&list=PLS--ClexQbQ1lg6TttQTcE3a60T_noSnh)). The series depicted the romantic relationships and friendships among six fictional university students, with dynamic changes in their relationships over time, such as characters becoming romantically involved, breaking up, experiencing love triangles, and unrequited love. None of the participants had prior exposure to this series. Participants were instructed to pay close attention to the social interactions and relationship changes in the series. The entire series was divided into three episodes (three fMRI scan runs), each lasting approximately 16, 15, and 11 minutes. Ten- and sixteen-second blank screens were included at the beginning and end of each episode, respectively. To ensure a seamless viewing experience, 30-second scenes from the end of the previous episode were repeated at the beginning of the subsequent episode. Data from these repeated scenes were excluded from analysis.

#### Rating task

After movie-viewing, participants were asked to rate the relationships between characters and the personality traits of each character, using a 5-point Likert-type scale. Neuroimaging data obtained during the rating task were not analyzed in this study. This task involved six relationship statements, including friendship, love, trust, listening, duration of friendship, and perceived co-occurrence in the movie, and seven personality statements, including dominance, warmth, extraversion, trustworthiness, competence, openness, and attractiveness. Each rating trial began with a fixation point displayed at the center of the screen for 500 ms, followed by the presentation of the names and faces of two characters at the top center of the screen along with the corresponding relationship statement. For example, a statement “Character B trusts character C” was shown alongside the names and images of character B and C. Participants provided their ratings, indicating their level of agreement with the statement on a scale ranging from 1 (strongly disagree) to 5 (strongly agree). In cases where participants were unable to assess a statement due to insufficient information in the movie, they were instructed to respond with “I don’t know,” although they were encouraged to provide a rating whenever possible. The “I don’t know” response was treated as 0 in subsequent analyses. Each relationship statement was rated for every directional pair of characters (e.g., both “Character B trusts Character C” and “Character C trusts Character B”). The duration of each trial was dependent on the participant’s response time, with a maximum limit of 12 seconds for their responses, except for one participant who completed the rating task with a fixed duration of 6 seconds. Once a response was provided or if 12 seconds elapsed without a response, the ongoing trial automatically concluded, and the subsequent trial began. The average response time for relationship rating trials was 3.04 seconds (*SD* = 1.40 seconds).

### Imaging procedure

#### Acquisition

Neuroimaging data were collected at the Center for Neuroscience Imaging Research of the Institute for Basic Science using a Siemens MAGNETOM Prisma 3 Tesla MRI scanner equipped with a 64-channel head coil. Functional data were acquired through a T2*-weighted echo-planar imaging (EPI) sequence (TR = 1000 ms, TE = 30 ms, FOV = 240 mm, multiband factor = 3, in-plane acceleration factor (iPAT) = 2, voxel size = 3 mm isotropic, 48 slices providing whole-brain coverage). High-resolution anatomical images were obtained using a T1-weighted Magnetization Prepared Rapid Gradient Echo (MPRAGE) sequence (TR = 2200 ms, TE = 2.44 ms, FOV = 256 mm, voxel size = 1 mm isotropic).

#### Preprocessing

Neuroimaging data were preprocessed using fMRIPrep (*69*) (version 23.0.2) with default parameters, including corrections for head motion and spatial normalization by aligning each participant’s brain to the MNI152NLin2009cAsym template. Afterward, nuisance regressors (*70*) were removed from the data, including six motion parameters along with their temporal derivatives, global signal, framewise displacement estimates, six anatomical component correction (*71*) (aCompCor) components derived from fMRIPrep, and polynomial regressors up to the second order. The blood-oxygen-level-dependent (BOLD) signals were scaled and spatially smoothed using a Gaussian kernel (FWHM = 6 mm). Lastly, the data were z-scored separately for each run.

### Regions of interest (ROIs)

We defined ROIs using the Brainnetome atlas, which parcellates the whole brain into 246 distinct parcels (*72*) (**Fig. S2**). The selected ROIs included the ventromedial (vmPFC) and dorsomedial prefrontal cortex (dmPFC), precuneus (PREC), temporoparietal junction (TPJ), and superior temporal sulcus (STS), regions previously implicated in social information processing (*57*, *73–75*). Additionally, we included regions involved in sensory processing, such as the early and lateral visual cortex (EVC, LVC), fusiform gyrus (FFG), and superior temporal gyrus (STG). Although these ROIs were specified in advance, all our analyses were performed across the whole brain, and voxelwise results are presented throughout.

### Movie annotation

This annotation dataset was previously described and utilized in the earlier publication (*76*). It is important to note that all neuroimaging and behavioral data were collected from an independent cohort of participants.

#### Detailed annotation

A trained annotator, who did not participate in the fMRI experiment or event segmentation, annotated the movie stimuli at one-second intervals. This annotator devoted approximately 40 hours to the task, producing a highly detailed and consistent annotation of characters’ actions, emotions, and interactions. This comprehensive annotation included four types of detailed annotations: 1) identifying the characters involved in social interactions as senders and receivers of the interactions, 2) describing the actions performed by each character, 3) identifying the emotions experienced by each character, and 4) providing a scene description. For example, in a scene at a pub where character A talks to characters B and C, and character A laughs while character C shows an angry expression, the detailed annotation included the following details: character A was designated as the sender and engaged in talking happily, and characters B and C were identified as receivers; character C was designated as the sender and their emotion was angry; and the scene was described as ‘characters A, B, and C are talking in the pub.’ For consistency, synonyms for action and emotion words (e.g., cry and sob) were replaced with representative words (e.g., cry).

#### Event segmentation

Four independent annotators, who did not participate in the fMRI scan and had not previously seen the series, were instructed to segment the series into discrete events by marking start and end time points whenever they identified transitions in topic, location, time, character, or relationship dynamics. For each segmented event, annotators also assigned a descriptive title and provided brief justifications for the identified boundaries (*77*). Event boundaries selected for behavioral modeling were those that reached consensus among at least three of the four annotators, within a five-second interval.

### Feature extraction

We developed a dynamic social interaction model to quantify social information in the movie using detailed annotations. Our approach involved measuring directional co-occurrence and affective graphs among characters in the series.

#### Co-occurrence graph

For the co-occurrence graphs, we counted the moment-by-moment frequency of characters appearing together. If characters A (sender) and B (receiver) were both present in the annotation, we assigned a directional co-occurrence score of 1 from A to B at time t (co-occurrence_AB,t_ = 1). For characters who did not appear together at the same time, the directional co-occurrence scores (e.g., co-occurrence_BC,t_ and co-occurrence_CB,t_) were set to 0.

To compute the event-wise co-occurrence matrix for behavioral model fitting, for each event, we generated a binary matrix where a value of 1 indicated the presence of co-occurrence between a directional pair, and 0 indicated its absence.

#### Affective graph

We measured the moment-to-moment valence of interactions between characters by computing sentiment scores for action and emotion words in the annotation using the Korean Sentiment Word Dictionary (*29*) (KSWD), which ranges from −2 (strongly negative) to +2 (strongly positive). The KSWD classifies meanings associated with approximately 12,000 words in the standard Korean dictionary into either positive or negative sentiments through a sentiment classification model. For each TR, we averaged these sentiment scores for all action and emotion words. Affective valence measures were defined only in cases where co-occurrence was detected, such that valence represents the qualitative aspect of an interaction conditional on its presence. For example, if character B was irritated and angry at time *t*, we assigned a sentiment score of −2 for the directional relationship between characters B and A (affective valence_BA,t_ = −2), based on the sentiment scores of the words ‘angry’ (−2) and ‘be irritated’ (−2) from the KSWD.

Next, we categorized the affective valence features into three groups: positive, negative, and neutral. Affective features with values in the top 2/3 of positive scores were classified as positive valence, while those in the bottom 2/3 of negative scores were classified as negative valence. Values that fell between these two ranges (−1/3 to 1/3) were classified as neutral valence. This approach allowed us to determine the affective valence of interactions between co-occurring characters, capturing not only their presence but also the nature and intensity of their interactions.

#### LLM evaluation

To validate the human-annotated interaction features and address potential annotation biases, we implemented an automated feature extraction pipeline using an LLM (Google Gemini). The LLM was prompted to act as an objective behavioral analyst and was provided with second-by-second character dialogues and scene descriptions, consistent with the human annotations. For each second, the model was instructed to extract two features: (1) the directionality of the social interaction, explicitly identifying the initiating (source) and receiving (target) characters, and (2) the affective valence of the source character, quantified on a 5-point scale from -2 (highly negative) to +2 (highly positive), with 0 indicating no affective valence. To ensure analytical consistency, interaction identification was restricted to the six main characters of interest.

### Behavioral modeling

For each event (*77*, *78*), we computed four character-by-character matrices representing co-occurrence, positive valence, neutral valence, and negative valence. The co-occurrence graph matrix was constructed by binarizing the presence of directional interactions between characters. For the affective graph matrices, we first aggregated sentiment scores for each event. These aggregated scores were then categorized into three valence groups using the same thresholding criteria described in **Feature extraction**: Interactions were categorized as positive, negative, or neutral based on percentile thresholds of the non-zero valence-score distribution. Positive interactions were defined as scores in the upper two-thirds of the positive range, negative interactions as scores in the lower two-thirds of the negative range, and neutral interactions as scores falling within the central range around zero. Based on this classification, we constructed three separate matrices. For the positive valence matrix, a value of 1 was assigned if character A directed a positive interaction toward character B, and 0 otherwise. For the negative valence matrix, a value of 1 was assigned for negative interactions, and 0 otherwise. For the neutral valence matrix, a value of 1 was assigned if the interaction was neither positive nor negative but still involved directed co-appearance, and 0 otherwise. This thresholding was applied to prevent any single event from exerting a dominant influence on model fitting. These matrices were summed across events and z-scored across character pairs to serve as model inputs. In other words, we generated one co-occurrence matrix and three valence matrices that captured directional social relationships across the entire series. To characterize how observing dynamic social interactions shapes perceived interpersonal relationships, we implemented a partial least squares (PLS) regression model to predict participants’ z-scored directional relationship ratings across four dimensions. The model was constructed using both co-occurrence and affective valence matrices as predictors (full model). PLS regression projected these matrices into a shared latent space while maximizing the covariance between the co-occurrence and affective feature sets and behavioral ratings.

To evaluate the contribution of each feature space, we performed a feature ablation analysis. Specifically, after fitting the full model with cross-validation, we set the weights associated with either the co-occurrence graph features or the affective graph features to zero, thereby removing their influence while leaving the latent structure of the model unchanged. This allowed us to compute prediction performance with one feature set (i.e., either co-occurrence or valence) effectively removed, and to compare it against the performance of the full model. By contrasting the reduction in prediction accuracy across these conditions, we quantified the relative importance of co-occurrence and affective features for reconstructing participants’ social relationship graphs.

#### Model evaluation

To assess model performance, we employed a leave-one-participant-out cross-validation (LOOCV) scheme. In each fold, beta coefficient matrices were estimated using data from 27 participants, and prediction accuracy was evaluated on the held-out participant by computing the R-squared value, resulting in a total of 28 folds. To generate null distributions for statistical testing, we applied an interaction-shuffling procedure. At each second, the identities of the interacting characters were randomly shuffled, and the event-level co-occurrence and three affective valence matrices were then reconstructed. The identical PLS regression and LOOCV pipeline was then applied to these shuffled features. This procedure was repeated 1,000 times, generating a null distribution against which observed R-squared values were compared to compute *p*-values.

#### Social relationship plot

To visualize latent representations of the PLS regression model (or any social relationship representations), we first scaled values in the representations through min-max normalization (0.001 to 1) and calculated the two-dimensional coordinates for each character using multidimensional scaling (MDS). We then aligned the shapes of the social relationship graphs using the Procrustes algorithm, adjusting the center, normalizing the spatial coordinates, and rotating them if needed.

#### Graph reciprocity

Reciprocity is a crucial aspect of social relationships (*32*). In social networks, reciprocity provides a measure of the directionality of relationships, such as unrequited love. We measured directed and weighted reciprocity following the method of the previous work (*79*). The reciprocated weight between two characters is defined as the minimum of the weights of the directed relationships between them. The non-reciprocated weight from character A to B is defined as the difference between the directed weight from A to B and the reciprocated weight. Based on these dyadic measurements, the reciprocated strength of a character is defined as the weighted sum of their reciprocated weights from others. Finally, the global reciprocity of the graph is measured as the ratio of the total reciprocated weight to the total weight. This measure provides an overall indication of how reciprocated the relationships in the network are.

### Voxelwise encoding modeling

#### Model training

Voxelwise encoding modeling predicts the BOLD responses of individual voxels based on features of the presented stimuli. This approach has been widely used to identify neural responses involved in processing visual (*80*, *81*) and semantic information (*82*), as well as in observing social interactions (*12*, *83*).

In our study, we used detailed annotations to extract moment-to-moment social information from the series, including co-occurrence and affective graphs, which collectively represent 30 directional relationships among six characters. To characterize the nature of social information encoded in each voxel, we built a voxelwise encoding model that predicts voxels’ BOLD responses using these two sets of features. We temporally smoothed this information using Gaussian filtering (σ = 5). To account for the hemodynamic response, all information regressors were convolved with the canonical hemodynamic response function (HRF). To confirm the robustness of the neural encoding model, we tested two additional variants of the model that controlled for lower-level visual signals (*81*) (motion energy), that did not apply temporal smoothing to social interaction information, and that applied FIR model instead of considering HRF convolution (**Fig. S3**).

For model training and validation, we used 6-fold cross-validation over the concatenated BOLD time series across each participant’s movie-viewing runs. As the stimulus consisted of three distinct episodes, each episode was split into two consecutive segments to generate six folds (yielding lengths of 493, 493, 457, 457, 333, and 333 TRs), with model weights estimated on five folds and evaluated on the held-out fold.

We employed banded ridge regression (*84*), an algorithm that incorporates regularization hyperparameters to effectively penalize non-predictive and redundant information. Unlike traditional ridge regression, prediction scores were obtained separately for each distinct type of information as well as for the entire set of information. We considered two information subspaces: one for co-occurrence (number of features = 30) and one for affective graphs (number of features = 30 x 3 = 90). The ridge regularization hyperparameters were optimized through a random search within a range from 10^0^ to 10^20^, with 41 log-scaled steps. This optimization process was performed separately for each type of information through a nested 5-fold cross-validation on the training data.

#### Model Performance

To assess the prediction accuracy of each model, we calculated Pearson correlations between the predicted voxel responses and the actual voxel responses on each test fold. These scores were Fisher Z-transformed and averaged across cross-validation folds within each participant. To assess the statistical reliability of average model performance at each voxel, we performed a hypothesis test using the one-sample bootstrap method (*85*). For each voxel, we built a bootstrapped distribution of average model performance by randomly sampling with replacement across participants and computing a new average performance score. This bootstrap procedure was repeated 5,000 times. To construct a null distribution, we centered the bootstrap distribution at zero by subtracting the observed mean from each bootstrap sample, and computed the one-tailed p-value as the proportion of this null distribution that was greater than or equal to the observed mean performance. To correct for multiple comparisons across voxels, we employed false discovery rate (FDR) correction using the Benjamini-Hochberg procedure (*q* < .05) for this and all subsequent analyses.

To estimate the individual contribution of each subspace, we used the split prediction score framework (*84*), which decomposes the total Pearson correlation between predicted and true responses into additive components attributable to each subspace. This split score involves predicting the target variable using the full model and isolating the contribution of each subspace by computing its marginal effect while controlling for the presence of the others.

#### Whole-brain plot

The Pycortex Python package (*86*) was used for visualizing the whole-brain results on the cortical surface.

### Reconstructing social relationship graphs

#### Neural reconstruction

To reconstruct participants’ internal representations of social relationships from neural activity, we first estimated voxelwise model weights for each type of social information using a banded ridge regression model. For each voxel, weights were averaged across six cross-validation folds, resulting in four weight matrices per participant: one for co-occurrence graphs and three for affective graphs (positive, neutral, and negative valence). Using these voxel weight matrices as predictors, we trained PLS regression models to reconstruct participants’ four-dimensional social relationship ratings, following the same procedure as in the behavioral modeling. All variables were z-scored prior to modeling, and model performance was evaluated using a LOOCV scheme. Reconstruction accuracy was quantified by computing Pearson correlations between the predicted and actual social relationship ratings for each participant. These correlations were Fisher Z-transformed for statistical testing. Voxelwise statistical significance was assessed using one-sided one-sample t-tests, and multiple comparisons were corrected using the Benjamini-Hochberg FDR correction procedure. For visualization, the weighted sum of the latent representations, combined according to their explained variance ratios, was projected onto a two-dimensional space using multidimensional scaling.

#### Dimensionality of relationship representations

To evaluate how the complexity of social relationship representations varied across the brain, we estimated the number of latent dimensions needed to reconstruct social relationship graphs from neural data. Specifically, for each voxel that showed statistically significant reconstruction similarity, we identified the minimum number of latent dimensions required to achieve 95% of its maximum possible reconstruction similarity. We implemented PLS regression models using one to four latent dimensions (the maximum number of latent dimensions allowed when using co-occurrence and three affective valence weight matrices). For example, if a voxel reached 95% of its maximum reconstruction accuracy using a single latent dimension, its dimensionality was recorded as 1; if two latent dimensions were needed, the dimensionality was recorded as 2, and so on. This analysis was conducted for each participant, and the resulting dimensionality values were averaged across participants at the voxel level. Finally, we visualized the dimensionality map on the cortical surface to examine regional differences in representational complexity across the brain.

### Shared neural spaces for social graph encoding

#### Shared Response Model (SRM)

To interpret neural representations of social graphs, we identified low-dimensional neural spaces using the SRM (*87*). SRM was applied to the voxel weights for social graph features (30 relationship pairs x 4 graphs = 120 features) derived from the voxelwise encoding model, identifying a common representational space for social graph encoding across participants. This approach reduces noise and dimensionality while preserving information shared among participants.

For each participant *i*, the data matrix X*_i_* ∈ R^VxF^ (voxels × features) was decomposed as X_i_ ≈ W_i_S, where W_i_ ∈ R^V×k^ (voxels × components) is the participant-specific transformation matrix and S ∈ R^k×F^ (components x features) is the shared component matrix. The number of shared components (*k*) was determined using LOOCV: SRM was trained on N−1 participants, the held-out participant was aligned to the resulting common space, and reconstruction accuracy was quantified as the correlation between the original and reconstructed voxel patterns. Peak reconstruction accuracy was observed at *k* = 3 (**Fig. S7A**).

The transformation matrices W_i_ provided voxelwise loadings for each shared component. To visualize their spatial distribution, voxel loadings were z-scored within each participant and averaged across participants. The resulting brain maps illustrate how different cortical systems contribute to each shared component. Note that positive and negative values should be interpreted as opposite directions within the same representational dimension, not as differences in activation magnitude.

The transformed graph features (30 relationship pairs × 4 graphs) for each of the three shared dimensions were reshaped into 6 × 6 character-by-character adjacency matrices (self-loop values set to zero) and used to predict four-dimensional relationship ratings in the PLS regression models. Graph reconstruction of social relationships for each component was performed by taking the weighted sum of the X-side latent representations, combined according to their explained variance ratios. Each reconstructed graph was then scaled using min–max normalization to the range [0.001, 1]. Finally, we characterized the representational structure of the three reconstructed social graphs using standard graph-theoretic measures.

#### Graph-level analysis

To characterize the structure of the social graph derived from the SRM, we applied five complementary graph-theoretic analyses (*88*).

1. In-going and out-going strength. For a directed weighted adjacency matrix *A* representing the strength of social relationships among six characters, each element *A*_ij_ denotes the directed weight from character *i* to character *j*. For each node, total incoming (in-going) and outgoing (out-going) connection strengths were computed as the sum of all corresponding edge weights.
2. Strength centrality. Strength centrality was defined as the mean of the in-going and out-going strengths for each node. This measure reflects the overall level of closeness of each character within a shared component.
3. In-out edge strength ratio. To capture the directional balance of connections, we calculated the natural logarithm of the ratio between in-going and out-going strengths. Positive values indicate that a node receives stronger inputs than it sends out, negative values indicate the opposite, and values near zero denote a balanced exchange of connection strengths.
4. Node reciprocity. For each pair of nodes, we identified the smaller value between the directed weights *A*_ij_ and *A*_ji_, representing the mutually shared portion of their connection (*79*). Summing these minimum values across all pairs yielded each node’s total reciprocal strength. Node reciprocity was then computed as twice this reciprocal strength divided by the sum of all in-going and out-going strengths of the node. High reciprocity values indicate that most of a node’s relationships are bidirectional and similar in strength, whereas low values reflect predominantly one-sided connections.
5. Local clustering coefficient. We defined a node’s local clustering coefficient by computing how densely its directly connected neighbors (those that either send to or receive from the node) are interlinked with each other (*89*). In the weighted, directed graph, we first normalized edge strengths to the unit interval and used a geometric-mean scheme to weight triangles: specifically, we took the cube root of the normalized weights and aggregated over triads so that stronger and more reciprocated links contribute more. Directionality was incorporated by symmetrizing these cube-rooted weights across both directions before counting weighted triangles. The coefficient is then obtained by dividing the node’s weighted triangle count by the number of possible neighbor–neighbor connections for a directed graph with reciprocated pairs properly accounted for; values are set to zero when this denominator is non-positive. High coefficients indicate that a node’s neighbors form a tightly knit subgroup, whereas low coefficients indicate that the node connects otherwise weakly linked or disconnected neighbors.

### Predicting trait impressions

Using a ridge regression model, we tested whether these relational graph metrics could predict perceived personality traits of the movie characters (e.g., dominance, warmth, extraversion, competence, trustworthiness, openness, and attractiveness). Relational features (input) and personality traits (output) were z-scored within participants. We trained the model on N - 1 participants and tested it on the remaining participant.

## Supplementary Materials

**Fig. S1.**
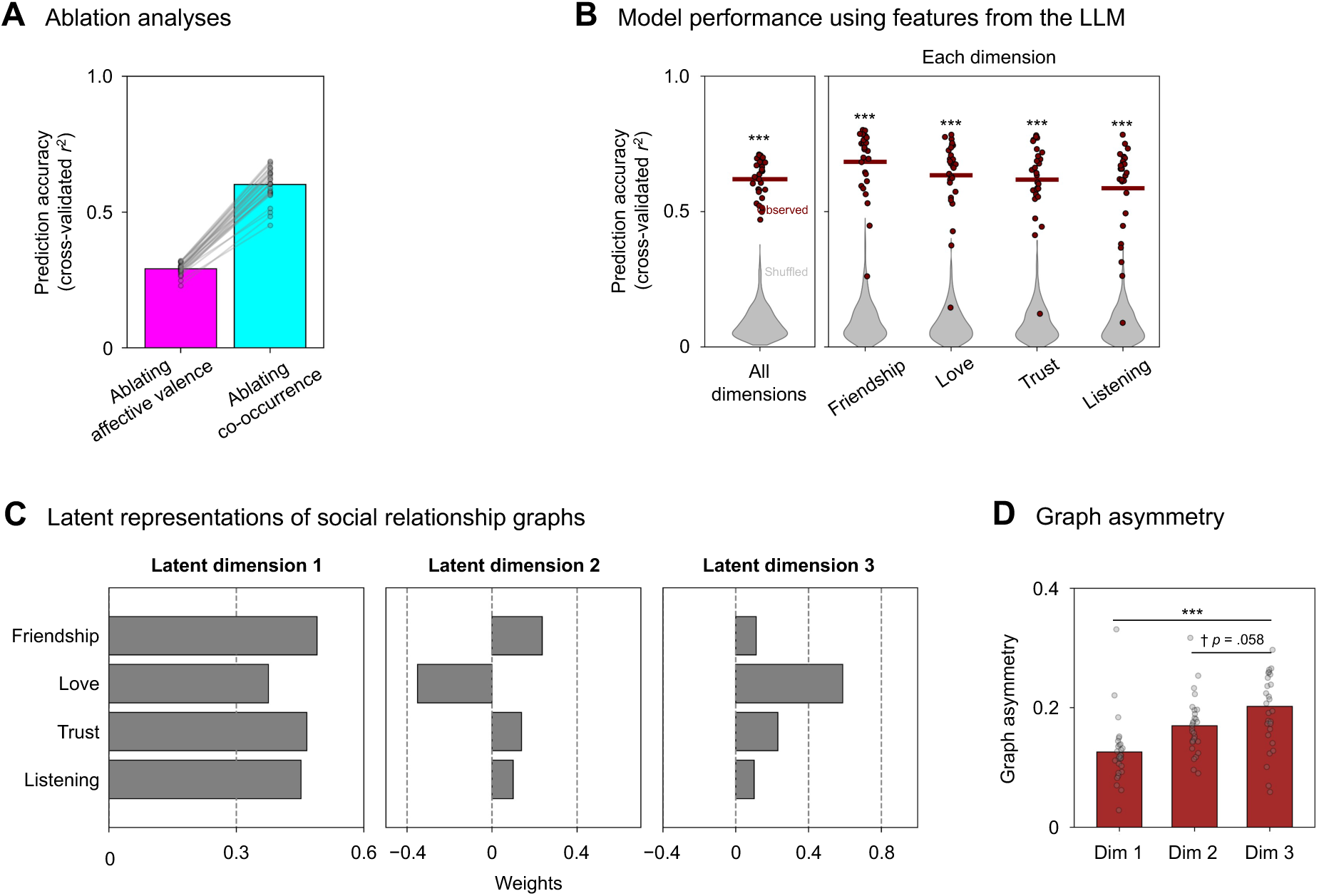
(**A**) Zeroing-out prediction accuracy was computed separately for co-occurrence and affective valence features. (**B**) Cross-validated prediction accuracy was computed by using interactional features from the LLM. To generate null distributions, we randomly shuffled the identities of interacting characters in annotations at each second, repeating this process 1,000 times. (**C**) Weight scores indicate how each relationship rating contributed to each latent dimension of the full model combining co-occurrence and affective graph features. (**D**) Graph asymmetry was quantified as 1 − graph reciprocity. The third dimension from the model showed higher asymmetry compared to the first and second dimensions. *** *p* < .001.

**Fig. S2.**
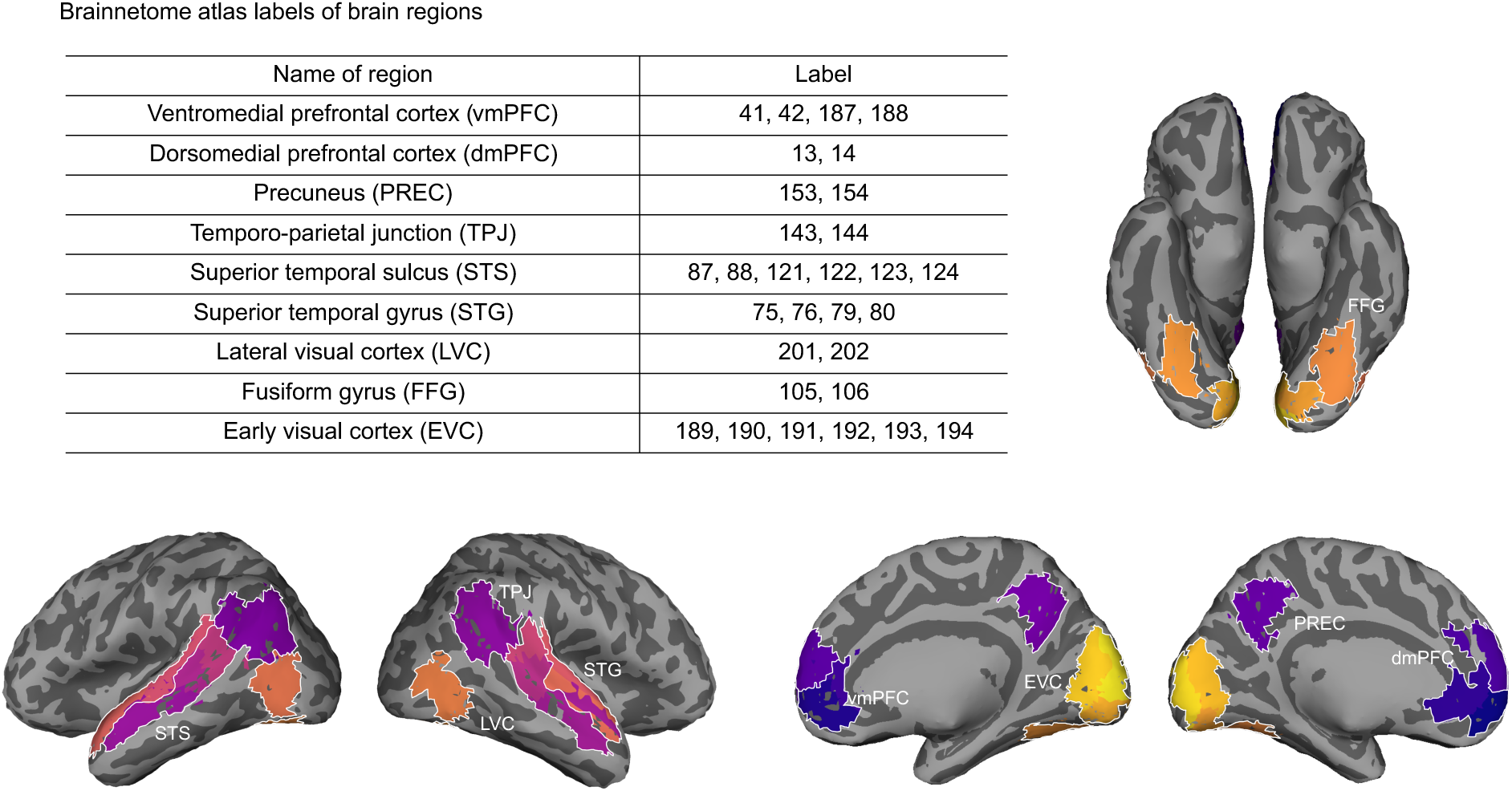
Label numbers of brain regions in our analysis and spatial locations of those brain regions.

**Fig. S3.**
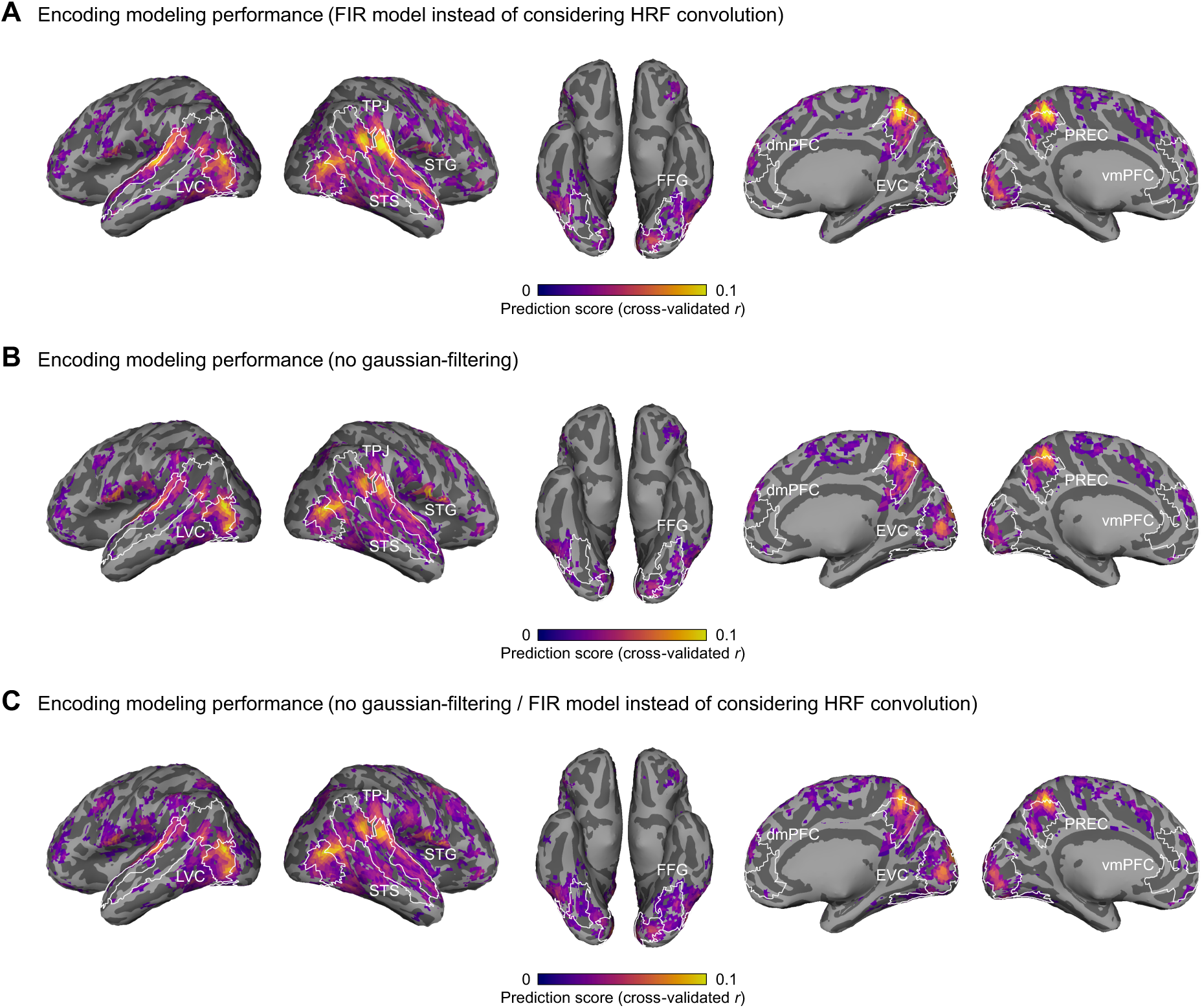
Prediction score maps of neural encoding models under different settings: (**A**) utilizing FIR model instead of considering HRF convolution, (**B**) utilizing social graph information without applying Gaussian filtering, and (**C**) utilizing FIR model instead of considering HRF convolution and social graph information without applying Gaussian filtering (*q*s < .05, FDR correction, clustering threshold = 20 voxels).

**Fig. S4.**
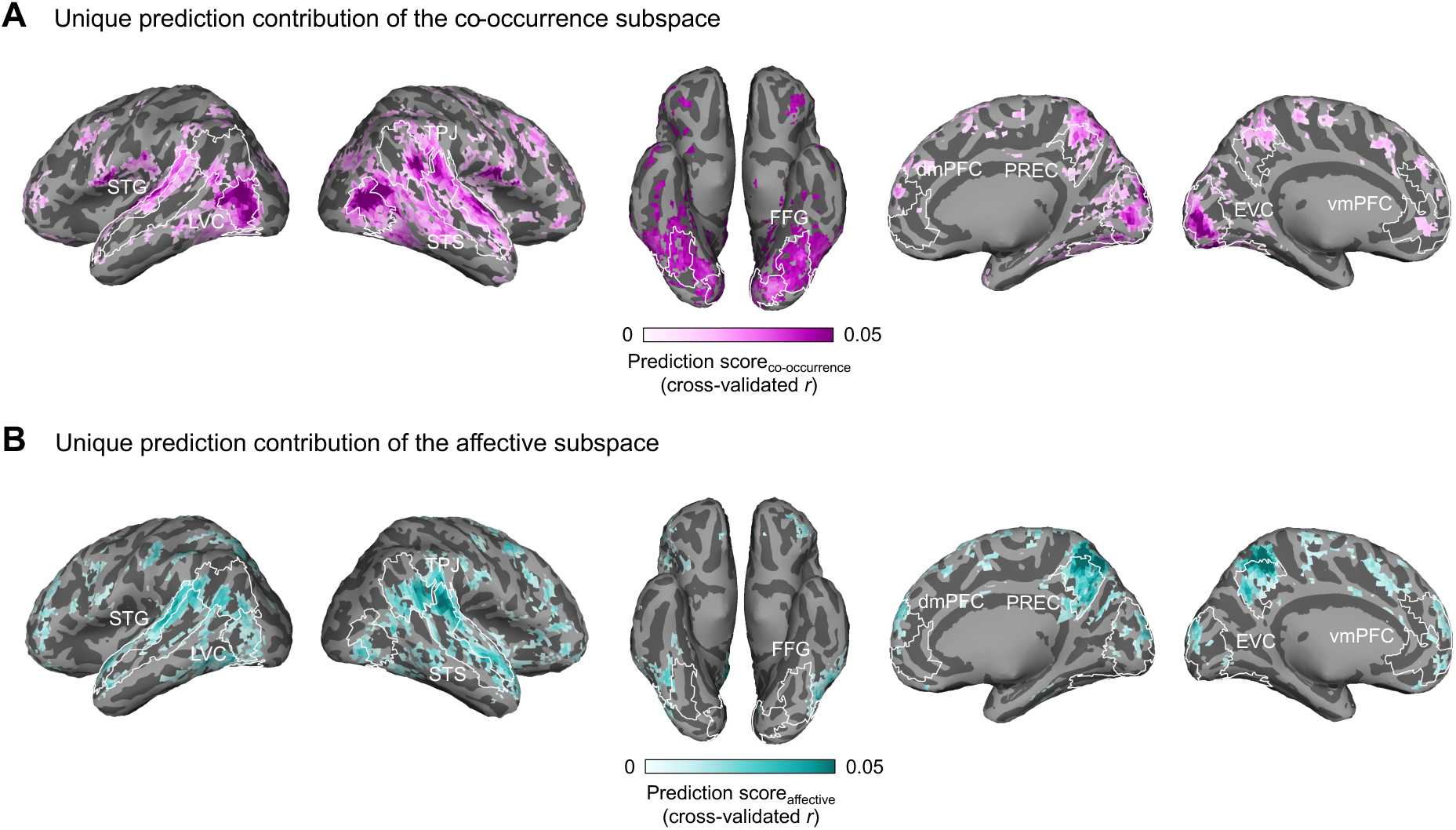
Only voxels with significant unique contributions from each graph are shown (for co-occurrence graph (**A**): *p*s < .05, clustering threshold = 20 voxels; for affective valence graph (**B**): *p*s < .05, clustering threshold = 20 voxels).

**Fig. S5.**
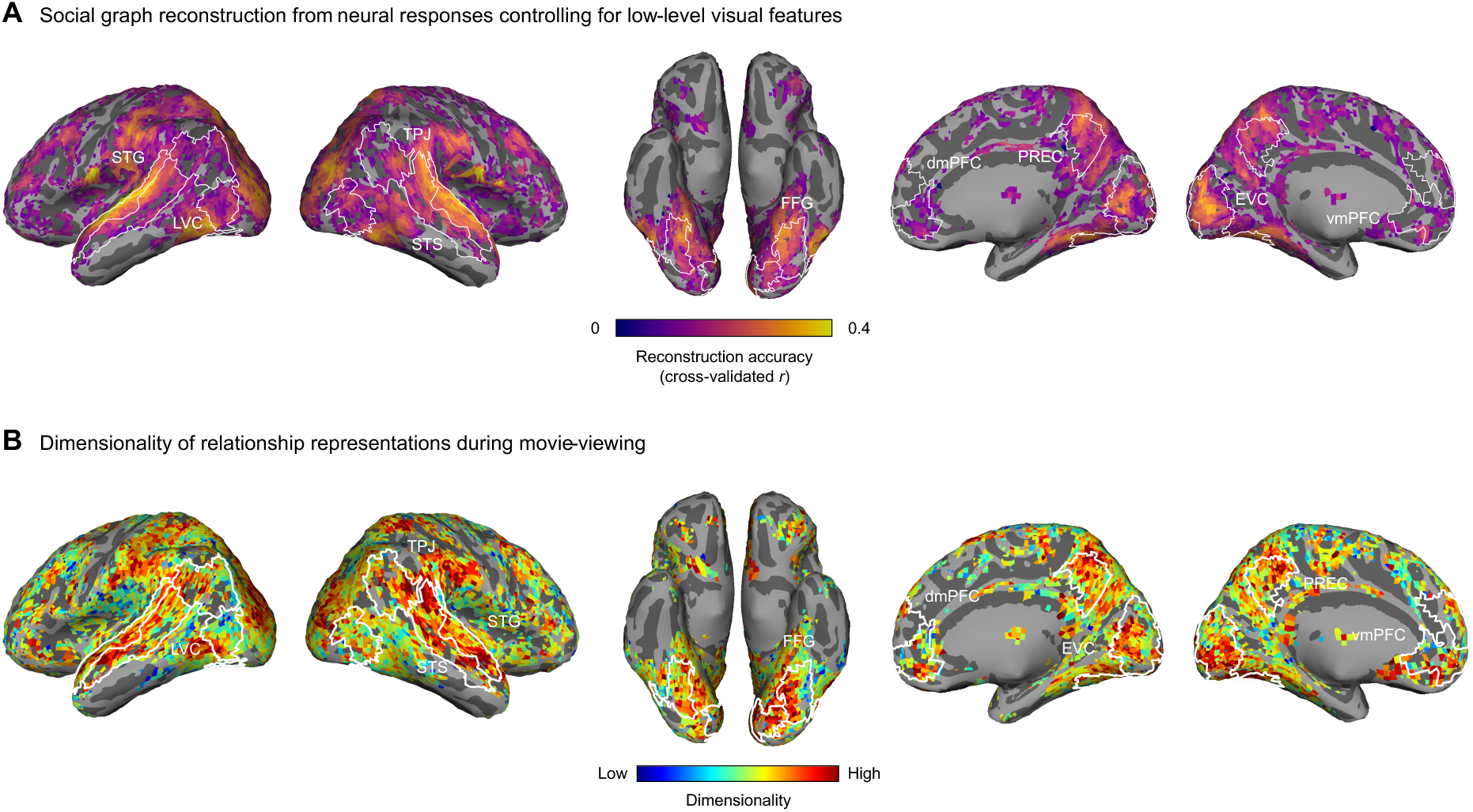
(**A**) Social relationship graphs were reconstructed from voxelwise encoding model weights (controlling for low-level visual features) using PLS regression with a LOOCV scheme. Reconstruction accuracy was quantified as the Pearson correlation between the predicted and actual four-dimensional relationship ratings. Only voxels with significant reconstruction accuracy are shown (*q*s < .05, FDR correction, clustering threshold = 20 voxels). (**B**) The dimensionality of social relationship representations was estimated for significant voxels in neural reconstructions by counting the number of latent dimensions required to explain 95% of maximum reconstruction similarity.

**Fig. S6.**
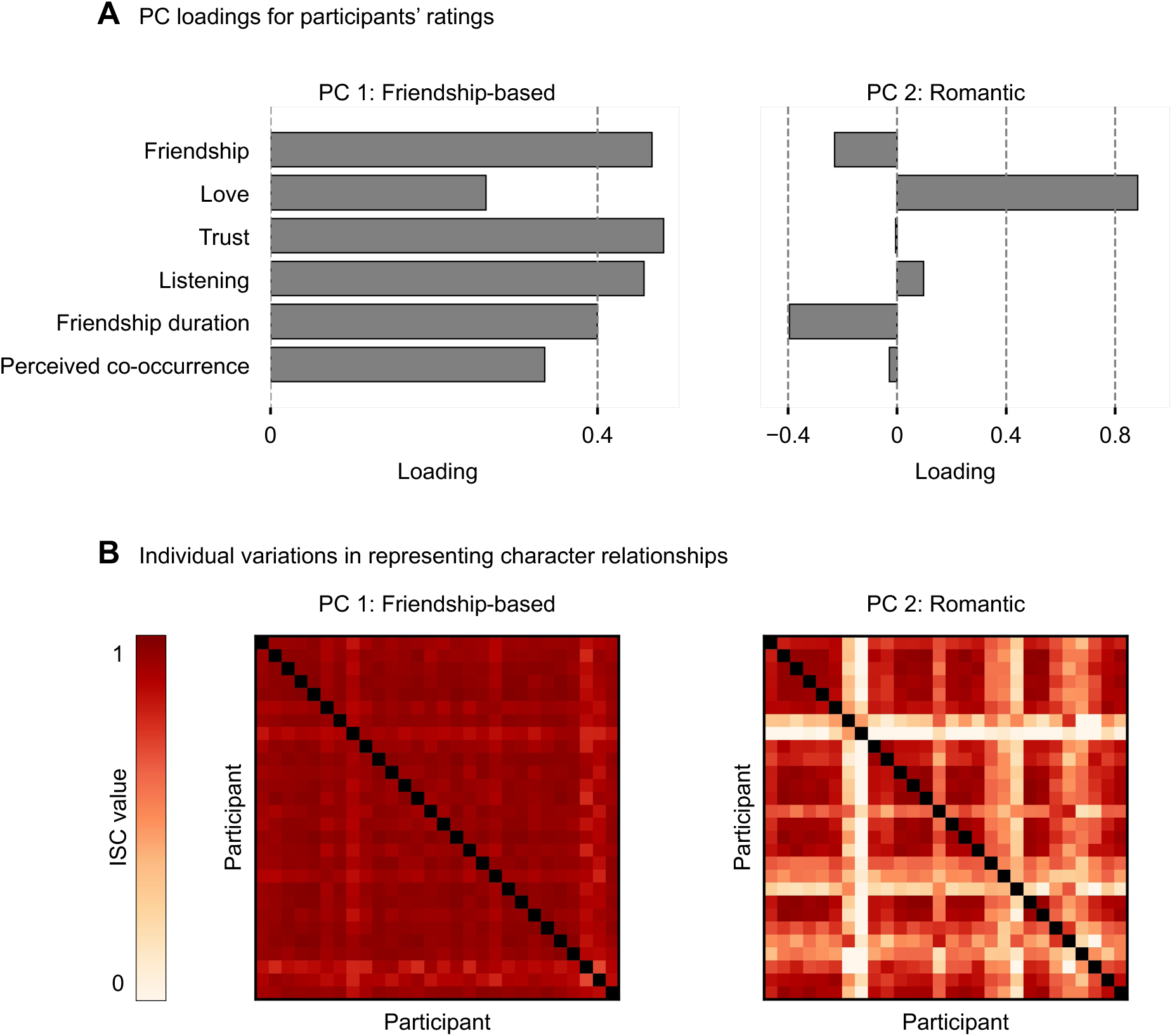
(**A**) The plot illustrates the loading scores for each relationship rating on Principal Component (PC) 1 and PC 2, derived from Principal Component Analysis (PCA). PC 1 exhibited high loadings on all relationship ratings except for love. Conversely, PC 2 had a high loading specifically on love but relatively lower loadings on the other ratings. Note that the sign of the loadings for PC 1 was reversed for the purposes of interpretation and analysis. (**B**) Inter-subject similarity matrices of character relationships based on the PC 1 and PC 2 estimation values. Participants’ responses varied more regarding the romantic relationship component (mean ISC value (r) = .62) than the friendship-based component (mean ISC value (r) = .93).

**Fig. S7.**
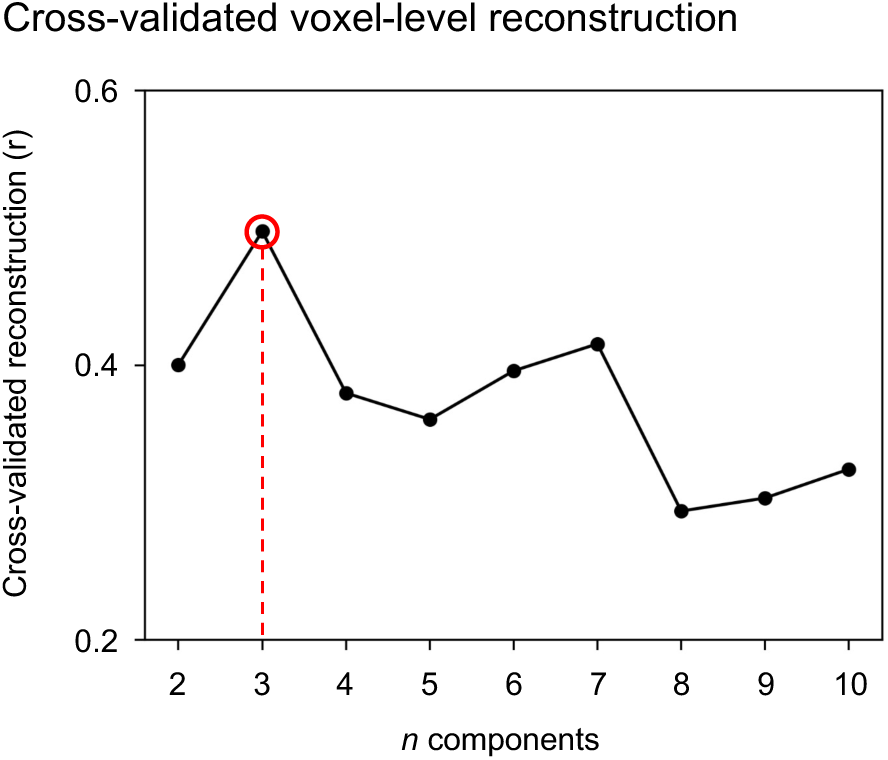
The optimal number of shared components for the SRM was determined using an LOOCV procedure. SRM was trained on 27 participants, and reconstruction accuracy of voxelwise neural weight patterns in the held-out participant was computed as the Pearson correlation between original and reconstructed patterns. Reconstruction accuracy peaked at k = 3, which was used for all SRM analyses.

**Table S1.** Rating task details. Participants rated six relationship dimensions among characters using a 5-point Likert scale and 0.

| Rating type | Statement |
| --- | --- |
| Friendship | A thinks B is close friend of A. |
| Love | A loves B. |
| Trust | A trusts B. |
| Listen | A listens B. |
| Friendship duration | A have known B a long time. |
| Perceived co-occurrence | A appeared together with B in the movie. |

**Table S1.** Rating task details. Participants rated six relationship dimensions among characters using a 1106 5-point Likert scale and 0.
| Rating score | Meaning |
| --- | --- |
| 1 | I strongly disagree with the statement. |
| 2 | I disagree with the statement. |
| 3 | Neither agree or disagree. |
| 4 | I agree with the statement. |
| 5 | I strongly agree with the statement. |
| 0 | I don't know. |

## Footnotes

1 Control analyses confirmed that the inclusion or exclusion of the two auxiliary dimensions (friendship duration and perceived co-occurrence) yielded qualitatively similar results.

## Notes

### Competing Interest Statement

The authors have declared no competing interest.

### Summary of Updates

The manuscript is revised. Introduction, Results, Discussion, Methods sections and Figures are revised.

